# Age-related biological dysregulation quantified using metabolomics is composed of distinct subsystems with synergistic associations to health outcomes: An analysis of the Canadian Longitudinal Study on Aging

**DOI:** 10.64898/2026.10.27.762256

**Authors:** Hesam Hafezalseheh, Michael Chong, Alan A. Cohen, Guillaume Paré, Andrew D. Rutenberg, Vanessa Taler, Dawn M.E. Bowdish, Chris P. Verschoor

**Author notes:** Corresponding author Chris Verschoor, MSc, PhD, Health Sciences North Research Institute, 56 Walford Rd, Sudbury, ON, Canada P3E 2H2.

## Abstract

Resilience is a fundamental component of healthy aging, whereby an organism can efficiently respond to external stressors and return to a resting biological and physiological state, known as homeostasis. This capacity declines with age in a heterogenous manner and is aggravated by biological dysregulation, a detrimental state broadly associated with health risk factors and poor outcomes. However, the underlying structure of dysregulation remains unclear, particularly in the context of older adult health trajectories. Here, we take a novel systems-based approach to the quantification of dysregulation using metabolomic data from n=9,992 older adults enrolled in the Canadian Longitudinal Study on Aging. We show that dysregulation across different biological systems is heritable, and exhibit distinct associations with health risk factors, indicators and outcomes, and DNA methylation patterns. Importantly, the effect of systems-specific dysregulation is cumulative, resulting in a significantly higher risk of disease incidence and death in older adults. Our findings broaden our understanding of biological dysregulation as a health state and demonstrate the value of system-specific dysregulation for effective strategies in personalized medicine and risk stratification.

## Introduction

Resilience, one of the fundamental pillars of healthy aging, is the intrinsic ability of an organism to efficiently respond to external stressors and return to a resting biological and physiological state, known as homeostasis. It is widely accepted that resilience is compromised with aging^1^ and is believed to significantly contribute to age-related chronic conditions and the loss of functional capacity^2^. The processes underlying resilience can be defined as intricate, overlapping networks of cells, molecules, and organ systems, with the complexity of these networks increasing in relation to the number of biological and physiological systems involved^2^. These processes are often categorized into interconnected “hallmarks”, such as inflammatory regulation, nutrient sensing, and mitochondrial function, that underggo significant changes as individuals age^3^ and function to maintain homeostasis in a collective fashion^4^. When this interconnected network of systems begins to break down, the result is colloquially referred to as biological dysregulation^5^. Dysregulation represents instability and diminishes the effectiveness of a stress response.

As resilience at the organism level is known to breakdown with age, dysregulation may further exacerbate this process as well as accelerate with acute illness and chronic disease^6^. Dysregulation has been shown to be associated with health risk factors and predict adverse health outcomes, including all-cause mortality and comorbidities, such as kidney disease and cardiovascular disease^7–12^. While dysregulation within specific systems, such as lipid metabolism and oxygen transport, has been shown to evolve through similar yet parallel processes^8,10,12^, this pattern is likely to depend on the nature of the stressor. Given that dysregulation is increasingly recognized as an important measure for effective strategies in personalized medicine, particularly improved diagnostic accuracy and more tailored interventions, characterization of the system specific nature of dysregulation is critical. However, current studies have been limited by a relatively small biomarker pool, and overlapping biological and physiological assignment, which is generally insufficient to describe a complex system^8^.

One of the commonly used methods to quantify dysregulation is Mahalanobis distance (D_M_) pioneered by Cohen and colleagues^7,13^. Despite its widespread adoption, D_M_ is subject to a few statistical limitations, including the normality assumption and the infeasibility of computation for highly correlated biomarkers. Others have used network-based clustering methods to tease apart the underlying structure of biological dysregulation, although most have been limited to only a few dozen biomarkers, resulting in weakly informative clusters containing just one or two biomarkers^14,15^.

Identifying meaningful biological subsystems through clustering could be improved by using more comprehensive biomarker data, such as metabolomics. Metabolomics data capture a wider range of biological processes and may help reveal connections between multiple biological processes that form an individual subsystem^16^. In addition, metabolomics data have been shown to be informative predictors of mortality and comorbidities^17^.

In the following study, we aimed to characterize biological dysregulation both globally and at the level of specific subsystems using a panel of over 800 metabolites in nearly 10,000 participants of the Canadian Longitudinal Study on Aging (CLSA). Specifically, we employed a novel approach based on a mathematical metric called Wasserstein distance, which measures how different an individual biomarker profile is from a reference profile within a population and can handle highly correlated biomarkers. We hypothesized that certain biological processes are interconnected and that their dysregulation contributes unequally to the health outcomes. Lastly, using data from the UK Biobank we demonstrated that our method can be applied to other types of biomarkers while exhibiting similar associations with health risk factors and outcomes.

## Methods

### Study population

The Canadian Longitudinal Study on Aging (CLSA) is an ongoing population-based prospective cohort study whose design and methods have been previously described^18^. In this study, we used the comprehensive cohort, which includes 30,097 individuals aged 45-86 years at recruitment who underwent physical and cognitive tests at 11 data collection sites in seven Canadian provinces. Baseline data were collected from 2012 to 2015, with follow-ups every three years thereafter. In this study, we used baseline and first (2015-2018) and second (2018-2021) follow-up data. Risk factors and outcomes used in this study are summarized in Supplementary Tables 1 and 2. The analytical data include metabolomics, hematology and clinical biomarkers, genomics, and DNA methylation.

### Participant factors

Participants were characterized according to sociodemographic, lifestyle, dietary, health indicator, and inflammatory markers. Baseline records were used.

Sociodemographic factors included household income, education, and marital status. Lifestyle factors included physical activity, smoking status, and whether tobacco was consumed in the past 24 hours. Physical activity was assessed using the Physical Activity Scale for Elderly (PASE)^19^ and categorized into three groups^20^: sedentary (PASE≤40), light activity (40<PASE≤90), and moderate to intense activity (90<PASE).

Dietary factors included whether caffeine was consumed in the past 24 hours, and alcohol consumption frequency.

Health indicators included body mass index (BMI), frailty index (FI), nutritional risk, depression status, general health, life satisfaction, and distress level. Inflammatory biomarkers included tumor necrosis factor (TNF), interleukin-6 (IL-6), and C-reactive protein (CRP). BMI was categorized as underweight, normal weight, overweight, and obesity classes I, II, and III based on the World Health Organization (WHO) classification. The FI was derived from 76 deficits pertained to cognition, physical function, BMI category, health and chronic conditions, depression, self-rated health, and activities of daily living, as described by Kanters et al^21^. The FI was categorized into three groups: low (FI≤0.1), medium (0.1<FI≤0.2), and high (0.2<FI)^22^. The nutritional risk score was derived using the eight-item AB-SCREEN II questionnaire^23^ and categorized into tertiles as low, medium, and high nutritional risk. Depression was defined using the CESD-10, a 10-item questionnaire measuring the frequency of depressive symptoms, including feelings of depression, loneliness, hopefulness for the future, and restless sleep. The responses of the CESD-10 range from 0–4, indicating the severity of depression, and a cut-off of 10 or more is used to determine whether an individual has depression^24^. The Satisfaction with Life Scale (SLS)^25^ was used as a life satisfaction factor and categorized into: very dissatisfied (4.5<SLS≤9.5), dissatisfied (9.5<SLS≤14.5), average (14.5<SLS≤24.5), satisfied (24.5<SLS≤29.5), and very satisfied (29.5<SLS≤35.5). Distress was measured using Kessler 10-item Psychological Distress Scale (K10)^26^ and was categorized into: mild (9<K10≤19), moderate (19<K10≤24), severe (24<K10≤29), and very severe (29<K10).

### Hematology, immunology, and clinical chemistry biomarkers

Cytokines included in this study were C-reactive protein (CRP) measured using the Cobas 8000 modular analyzer (Roche Diagnostics), and TNF and IL6 measured using Quantikine high sensitivity ELISA (R & D Systems). Hematology biomarkers were measured using Coulter Ac·T diff Analyzer (Beckman Coulter) and included white blood cells (WBCs), granulocytes, monocytes, lymphocytes, red cell distribution width (RDW), hematocrit, hemoglobin, red blood cell count (RBC) and platelet count. Biochemistry biomarkers measured using the Cobas 8000 modular analyzer (Roche Diagnostics) included creatinine, free thyroxine (T4), thyroid-stimulating hormone (TSH), ferritin, triglycerides, low-density lipoprotein (LDL), high-density lipoprotein (HDL), cholesterol, non-HDL, albumin, alanine aminotransferase (ALT), hemoglobin A1c (HbA1C), N-terminal pro-brain natriuretic peptide (NT-proBNP), and troponin T. All biomarkers were measured at baseline, except for NT-proBNP and troponin T, which were measured at the first follow-up.

### Outcomes

Outcomes included cognitive and physical function, healthcare utilization, all-cause mortality and chronic conditions.

Cognitive function was assessed using normative cognitive scores and reliable cognitive change scores for memory and executive function^27^. These scores represent cognitive function at baseline and a three year change in cognitive function by removing cognitive variation due to language, education, age, and sex^27^. Cognitive tests were conducted in person at baseline and during the first follow-up but were administered by telephone during the second follow-up due to the COVID-19 pandemic. Therefore, we only considered the baseline and the first follow-up assessment for cognitive functions. Physical function tests included grip strength, measured using a wireless dynamometer, and walking speed (inverted walking time), measured using the 4-Metre Walk Test. Grip strength and walking time were collected in person across all study waves, including the second follow-up.

Healthcare utilization represents a participant emergency department visit or overnight hospitalization in the past 12 months of baseline data collection. Participant deaths were ascertained as of July 2015 until March 2022 through direct contact with the next of kin, linkage to provincial administrative databases, and interim follow-up calls conducted between the main data collection waves^28^. Chronic conditions were self-reported and included angina, chronic obstructive pulmonary disease (COPD), dementia and Alzheimer’s disease (AD), diabetes, heart attack, heart disease, hyperthyroidism, hypothyroidism, kidney disease, osteoarthritis, osteoporosis, Parkinson’s disease, rheumatoid arthritis (RA), and stroke. For incident diseases, participants who reported the disease at baseline were excluded from the analysis.

### Metabolomics

Metabolomic profiling was conducted on EDTA plasma samples collected at baseline from 9,992 randomly selected individuals using non-targeted liquid chromatography–mass spectrometry (LC-MS) (Metabolon, North Carolina, USA). A detailed description of sample handling, metabolomic profiling, and data processing has been previously deescribed^29^. A total of 1,458 metabolites were detected with 811 metabolites exhibiting less than 50% missing values in each batch were batch-normalized and imputed by the minimum value in each batch. The annotations of metabolites are available in Supplementary Table 4.

### External validity in the UK Biobank

The UK Biobank (UKB) is an ongoing population-based prospective cohort, including approximately 500,000 participants aged 37–73 years at recruitment between 2006 and 2010^30,31^. Extensive phenotypic data were collected through questionnaires, physical assessments, and biometrics.

Participants characteristics and risk factors included: frailty^32^, physical activity (i.e. metabolic equivalent of task (MET) as quantiles), household income, education, BMI, smoking, general health, and inflammatory markers, including TNF, IL-6, and CRP. Outcomes included 16-year all-cause mortality, and incident diseases, including all-cause dementia, angina, atrial fibrillation, COPD, diabetes, heart attack, heart failure, hyperthyroidism, hypothyroidism, kidney disease, osteoarthritis, osteoporosis, Parkinson’s disease, RA, and stroke. Participant deaths were ascertained as of July 2010 using linkage to the national death registry. Incident disease diagnoses were ascertained using the 10^th^ revision of the International Classification of Diseases (ICD-10) codes from self-reports, linked hospital inpatient records, primary care records, and death registry data.

To quantify dysregulation (see below), we employed both metabolomic and proteomic data. Metabolomic profiling was performed using the targeted Nightingale NMR platform for 275,241 participants. We conducted quality control by removing metabolites with more than 10% missing values across participants, resulting in 159 metabolites (Supplementary Table 5). Missing values were replaced with the minimum value for each metabolite. Proteomics data were generated using the Olink platform from blood plasma samples collected from 54,219 UKB participants^33^. After excluding proteins with more than 20% missing values and individuals with more than 50% missing proteins, 2,905 proteins and 50,575 individuals remained (Supplementary Table 6). Missing values were replaced with half of the minimum value for each protein after scaling the values within each batch.

### Quantifying biological dysregulation using the Wasserstein distance

Wasserstein distance is a mathematical metric used to quantify the difference between two probability distributions in terms of their mean, variance, and shape and is colloquially known as “earth mover’s distance”, the minimum amount of work required to transform a probability distribution to another. If biological dysregulation represents how different an individual biomarker profile is from a healthy reference profile, then the Wasserstein distance can be conceptualized as the minimum amount of work required to transform an unhealthy profile into a healthy profile. Given two continuous distributions *A* and *B* with mean *μ_A_* and *μ_B_*, variance *σ_A_* and *σ_B_*, and cumulative distribution function (CDF) *F_A_* and *F_B_*, the squared 2-Wasserstein distance can be expressed as^34,35^:

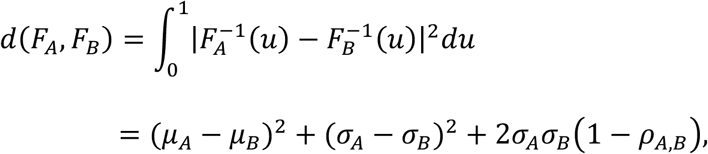

where 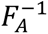 and 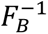 denote the quantile functions and *ρ_A_*_,*B*_ ∈ [0,1] is the Pearson correlation coefficient between the quantile-quantile (Q–Q) plot points corresponding to *F_A_* and *F_B_*. The third term of the formula, 2*σ_A_σ_B_*(1 − *ρ_A_*_,*B*_), captures the shape difference between the two distributions, including skewness and multimodality. Wasserstein distance was calculated using the R function ‘wasserstein_metric’ from the R package ‘waddR’^35^ v4.5. Biological dysregulation quantified using the Wasserstein distance was Box-Cox transformed and scaled to approximate a normal distribution. Individuals aged between 45 and 54 with a low frailty (FI<0.1) were selected as reference healthy population. The mean biomarker profile of the healthy population was then calculated and scaled with the rest of the population for each biomarker. To determine the age threshold of 54 years, we calculated the coefficient of variation in biological dysregulation based on metabolomics among the selected healthy reference population and identified the age at which the coefficient of variation was minimized. The use of age and FI thresholds reflect biomarker variations associated with age-related decline in resilience. The coefficient of variation in dysregulation among reference individuals aged 45-54 with FI<0.1 was the lowest compared with that among individuals of younger or older ages. The same approach was used for analyses of participants from the CLSA and UKB.

### Network analysis

To construct a metabolite network and cluster metabolites into biological subsystems, we used the unsupervised approach Weighted Gene Co-expression Network Analysis (WGCNA) from the R package ‘WGCNA’^36^ v1.73. To tune the WGCNA parameters, we used a combination of risk factors, health outcomes, and clinical biomarkers. We used the R function ‘blockwiseModules’ to build the network with multiple parameter combinations tested to maximize correlations between the first principal component (PC) score of system and health variables while minimizing the number of health variables associated with multiple systems PC score. The final parameters selected were: networkType = “signed”, TOMType = “signed 2”, soft threshold power= 7, minModuleSize= 40, deepSplit= 2, mergeCutHeight= 0.15, minKMEtoStay= 0.18, and pamRespectsDendro=FALSE.

To visualize the correlation network of metabolites, we used the R packages ‘igraph’ v2.1.4 and ‘plotly’ v4.11.0. To characterize the systems, we used LASSO regression with the PC score of each system as outcome and all health traits as predictors. A subset of traits was selected by LASSO as independent variables for predicting the PC score of each system. The LASSO regression coefficients were visualized using a heatmap generated with the R package ‘pheatmap’ v1.0.13. We performed pathway enrichment analysis for each system of metabolites using MetaboAnalyst 6.0^37^ with RaMP-DB database, which contains pathways from multiple resources, including HMDB, Reactome, WikiPathways, and PFOCR. Pathways with a Holm-adjusted p-value<0.01 and a minimum of five hits were determined to be statistically significant.

### Genetic and epigenetic analyses

To determine whether biological dysregulation is genetically determined, we estimated single nucleotide polymorphism (SNP)-based heritability using genomic data. The CLSA genomic data have been previously described^38^. In total, we included 8,299 unrelated individuals of European ancestry who had both metabolomics and genomics data available in the CLSA^39^. We followed the same genomic data preprocessing steps described in a previous publication using these data^39^. Briefly, we used Plink2^40^ to select autosomal SNPs with a minor allele frequency greater than 0.01, a missing rate less than 0.1, and an imputation quality control greater than 0.3. Additionally, we used UKB genomic data from 182,347 and 34,638 unrelated British White individuals who also had metabolomics and proteomics data, respectively. We estimated the heritability of biological dysregulation using the GCTA-GREML program^41^ v.1.94.1, adjusting for age, sex, genotyping batch, time since last meal or drink, and the first ten genetic PCs. To conduct epigenome-wide association analysis (EWAS), we used DNA methylation data from the CLSA, comprising 1,479 individuals with measurements at 862,927 cytosine-guanine dinucleotide (CpG) sites using the Illumina Infinium MethylationEPIC BeadChip microarrays on DNA extracted from peripheral blood mononuclear cells (PBMCs) samples at baseline. Outliers and low-quality samples were excluded as described by Verschoor et al^42^, resulting in a final sample of 1,446 participants for further analysis. We used the R package ‘minfi’ v.1.54.1 for preprocessing the data and extracting the beta values. We used the R package ‘limma’ v. 3.64.1 for performing EWAS on biological dysregulation adjusted for age, sex, and immune cell fraction. To estimate the immune cell fraction from the DNA methylation data, we used the R package ‘BloodCellDecon’ version v0.0.0.9000. CpG sites with p-values below 9.42×10⁻⁸, corresponding to a 5% family-wise error rate (FWER), were considered significantly associated with dysregulation^43^. To visualize the EWAS results, we used the R packages ‘ComplexHeatmap’ v. 2.24.1 and ‘Circlize’ v. 0.4.16.

Four epigenetic clocks from three different generations were selected: Horvath2013^44^, a first-generation clock trained to predict chronological age; GrimAge^45^ and PhenoAge^46^, second-generation clocks trained to predict mortality; and DunedinPACE^47^, a third-generation clock that measures the pace of aging and reflects longitudinal biological change.

### Statistical analysis

Associations between biological dysregulation and risk factors were examined using linear regression adjusted for age, with an interaction term between sex and each risk factor. To obtain sex-specific effects, we used the R function ‘emtrends’ from the R package ‘emmeans’ v.1.11.2. Biological dysregulation was regressed on epigenetic aging clocks using age-adjusted linear regression models with sex-by-clock interaction terms. To estimate sex-specific associations between overall dysregulation and outcomes, including cognitive function, physical function, healthcare utilization, and mortality risk, we used age-adjusted regression models with a sex-by-dysregulation interaction term. To estimate system-specific associations, we used regression models adjusted for age and sex. For all-cause mortality and incident disease predictions, we used Cox Proportional Hazards models, and participants were censored at their last data collection date or withdrawal date. We used linear mixed models with a participant-specific random intercept to assess physical function and a random effect for province to assess cognitive function. The interaction between dysregulation and wave of data collection represented changes in either cognitive function or physical function. For healthcare utilization outcomes, we used generalized linear models with a binomial family. The R packages ‘survival’ v.3.8.3 and ‘lme4’ v.1.1.37 were used to fit the Cox model and mixed models, respectively. We also categorized biological dysregulation into tertiles (low, medium, and high), and generated Kaplan-Meier curves using the R package ‘survminer’ v. 0.5.0. Regression results were reported in heatmaps and forest plots using the R packages ‘ggplot2’ v. 4.0.1, ‘ComplexHeatmap’ v. 2.24.1, and ‘forestploter’ v. 1.1.3. All statistical analyses were conducted using R v.4.5.0.

## Results

### Population characteristics

A graphical overview of our study design is presented in Fig. 1a. We included 9992 individuals with a median age of 62 years at baseline, of whom 51% were females and 96% were white. The household income of over 65% of the population is over 50,000 Canadian dollars (CAD) per year, with the average body mass index (BMI) and frailty index (FI) being approximately 28 (overweight) and 0.13 (not frail), respectively. Detailed characteristics of risk factors and chronic conditions are provided in Supplementary Tables 1 and 2.

**Fig. 1.**
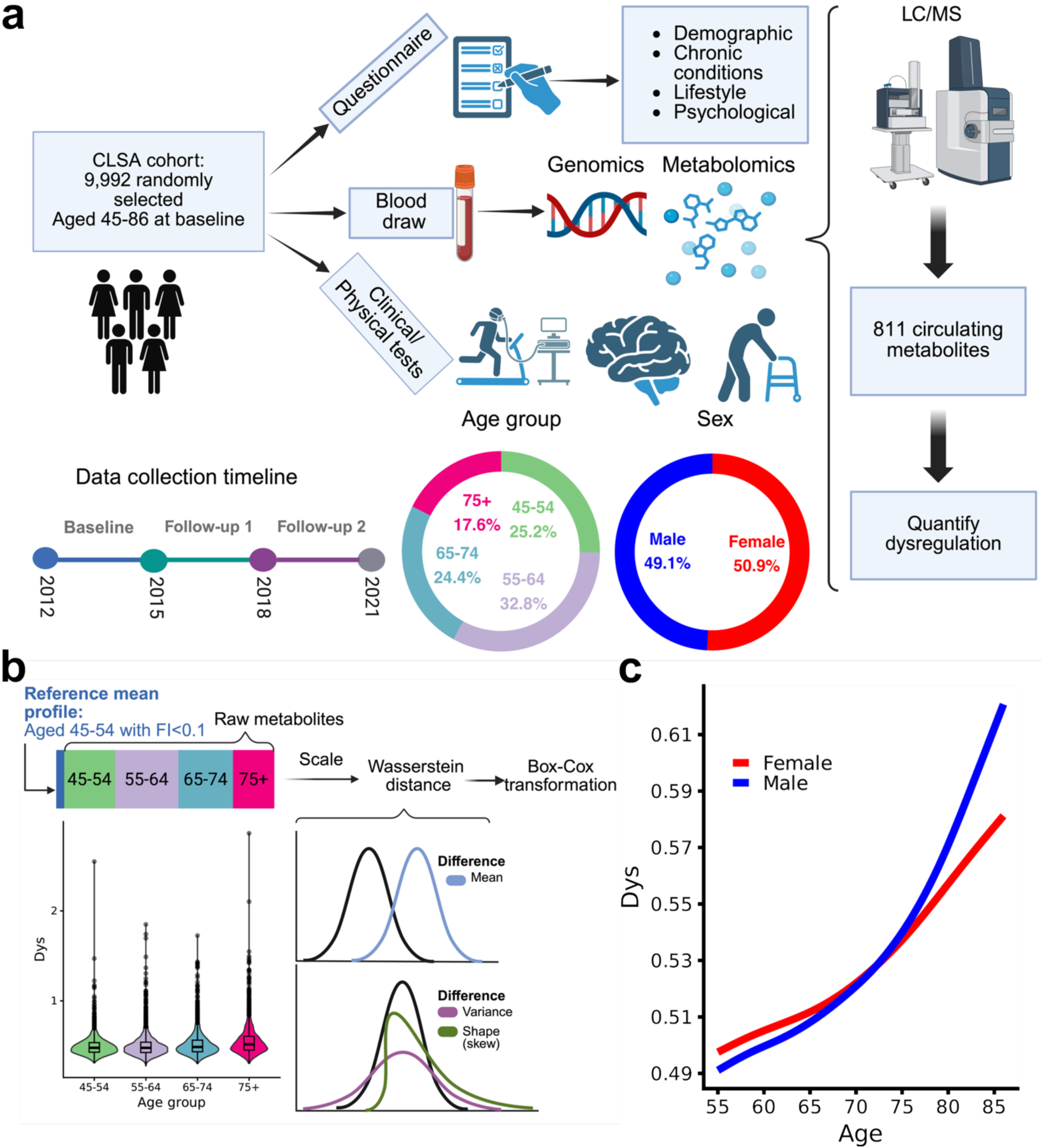
Study overview. **a**, Visual description of the Canadian Longitudinal Study of Aging cohort data employed. **b**, Workflow for quantifying biological dysregulation with a comparison of dysregulation (Dys) across age groups. **c**, The relative increase in biological dysregulation from age 55 to 86, shown separately for males and females.

### Quantifying biological dysregulation

We employed 811 metabolites to quantify biological dysregulation using the Wasserstein distance. Bivariate analysis indicated that the variance of dysregulation increases from younger to older age groups (p<0.001) (Fig. 1b), while the average magnitude increases with age in both males and females (p<0.001) (Fig. 1c). Although not statistically significant (p=0.3), males generally experienced a steeper increase in dysregulation with age compared to females. Dysregulation was also found to be heritable, where 21% (95% CI=8.6%-33.3%, p<0.001) of the variation in dysregulation can be explained by genetic factors. The remaining analyses included 7,471 individuals aged 55–86 years.

### Associations with risk factors and epigenetic age

We examined the associations of our novel dysregulation measure with a range of risk factors for males and females (Fig. 2a and 2b and Extended Data Fig. 1). We found that dysregulation was broadly associated with harmful sociodemographic, lifestyle, health indicators, dietary factors, and inflammatory biomarkers. In particular, dysregulation was significantly higher in participants characterized as: low-income earners (less than $20,000 per year; d=0.53), living a sedentary lifestyle (d=0.44), current, heavy smokers (d=0.51), underweight (d=0.70) or severely obese (d=0.39), highly frail (d=0.68), or under very severe distress (d=0.57). Dysregulation associations tended to be significantly higher in men than women, particularly for: high frailty (d_male_= 0.82, d_female_=0.55), living a sedentary lifestyle (d_male_=0.59, d_female_=0.28), and higher levels of inflammatory markers, including C-reactive protein (CRP), interleukin-6 (IL6), and tumour necrosis factor alpha (TNF) (d_TNF,male_=0.22, d_TNF,female_=0.14). Biological aging, as measured using four commonly used epigenetic clocks, was also associated with dysregulation (Fig. 2c). Particularly, dysregulation increased significantly with DunedinPACE, GrimAge, and PhenoAge, but not Horvath for both males and females (d= 0.03-0.19), with a sex difference observed for DunedinPACE (d_male_=0.19, d_female_=0.1).

**Fig. 2.**
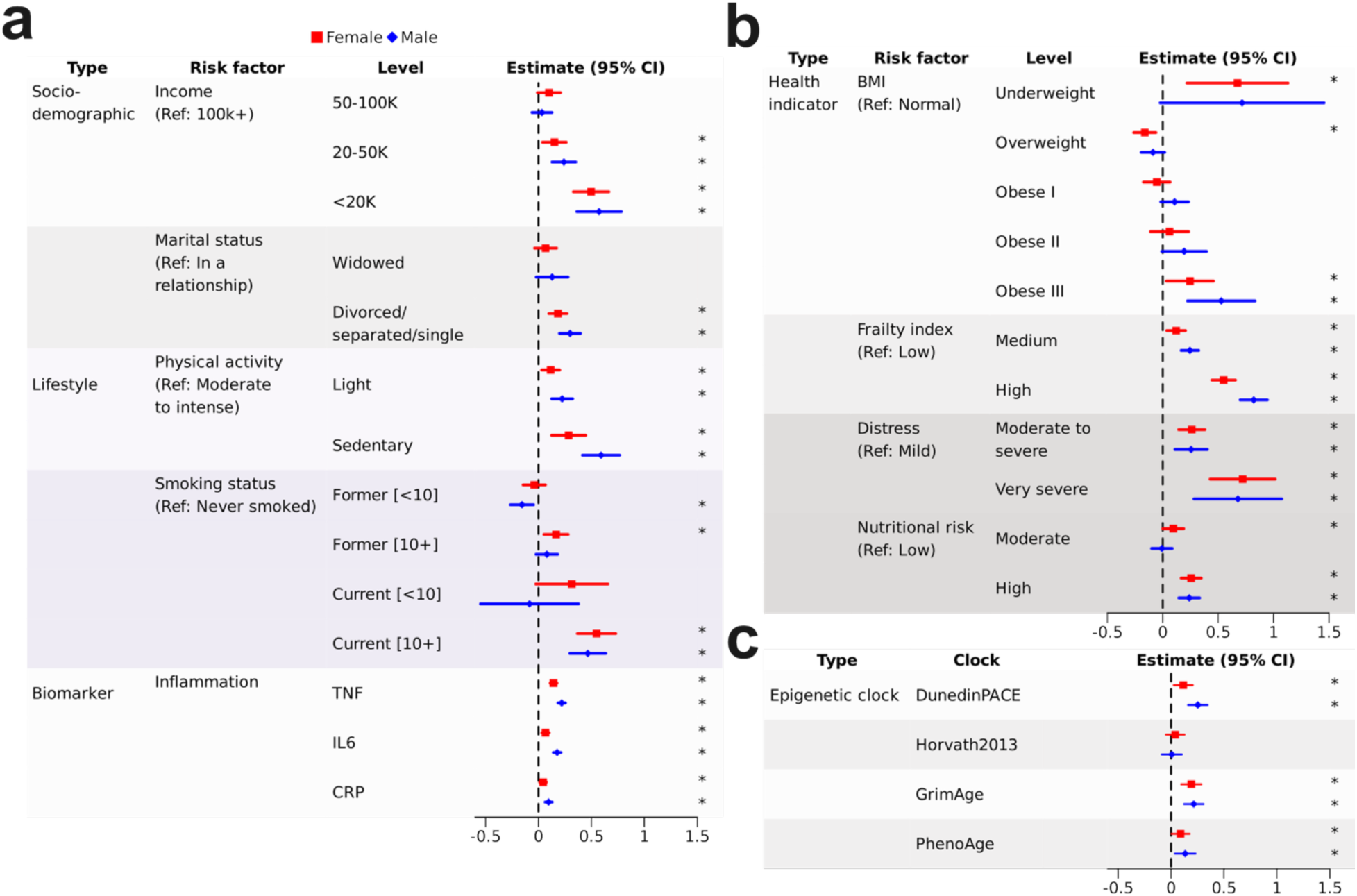
Sex-specific associations of dysregulation with health risk factors and epigenetic aging clocks. Standardized estimates and 95% confidence intervals from linear regression analyses describing the association between dysregulation and **a,** sociodemographic, lifestyle factors and inflammatory biomarkers, **b,** health indicators, and **c,** epigenetic clocks. All models include the fixed effects of age and the interaction between sex and a single predictor. The dashed line indicates no correlation, while the asterisks ‘*’ indicates p<0.05.

### Health outcomes associated with dysregulation

We also investigated whether dysregulation is predictive of health-related outcomes, including cognitive and physical function as well as healthcare utilization and nine-year all-cause mortality (Fig. 3 and Extended Data Fig. 2). Dysregulation was cross-sectionally associated with both memory and executive function for both males and females (d ranging between -0.06 and -0.05, Fig. 3a). Additionally, a 1-s.d. increase in dysregulation at baseline was associated with a small (d=-0.04), but significant decline in executive function among males at three years of follow-up. For physical function, we found dysregulation to be associated with weaker grip strength (d_male_= -0.09 and d_female_=-0.05, Fig. 3b) and slower walking speed (d_male_=-0.11 and d_female_=-0.09, Extended Data Fig. 2b) at baseline for both sexes, with a significant sex difference for grip strength. It was also associated with the decline in grip strength after six years, although only for males (d=-0.03). For both sexes, dysregulation was associated with higher odds of visiting an emergency department (Fig. 3c) or being hospitalized overnight (Extended Data Fig. 2b) in the last 12 months (ORs=1.21-1.39). Lastly, survival analyses showed that individuals with higher dysregulation have lower survival probabilities, where a 1-s.d. increase in dysregulation was associated with a 59% (CI=43%-78%) and 49% (CI=28%-74%) higher risk of mortality in males and females, respectively (Fig. 3d). While the sex difference in mortality risk was not statistically significant, we did observe a non-linear pattern in this relationship (Extended Data Fig. 2c). Notably, the risk of mortality due to increased dysregulation remained significant after additional adjustments for epigenetic age or frailty, suggesting that dysregulation has an independent effect from other health measures (Extended Data Fig. 2.d).

**Fig. 3.**
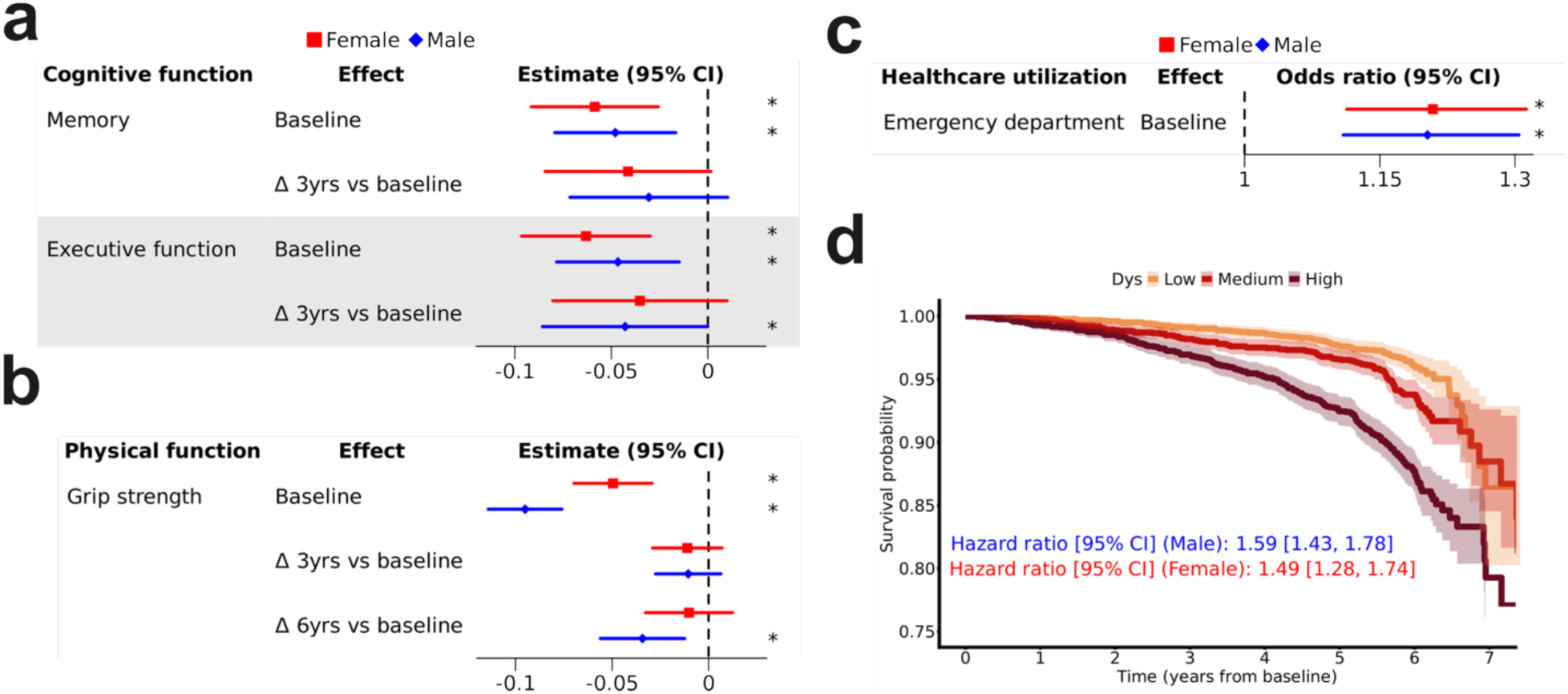
Sex-specific associations of dysregulation with functional decline and health outcomes and survival analysis. Standardized estimates and 95% confidence intervals reported for the association between dysregulation and **a,** memory and executive function, and **b,** grip strength. Δ 3yrs or Δ 6yrs vs. baseline refers to the 3-year or 6-year change relative to baseline, and significance denoted by asterisks (p<0.05). **c,** Odds ratios and 95% confidence intervals for emergency department visits associated with dysregulation. **d,** Associations with all-cause mortality indicated by Kaplan-Meier curves, with dysregulation tertilized as low, medium or high, and hazard ratio (95% confidence interval) estimates derived from Cox proportional hazards regression. All regression models included the fixed effects of age and a sex by dysregulation interaction term.

### External validation using data from the UK Biobank

To validate our novel approach and replicate our results in another cohort with other types of data, we used metabolomic (n=159 metabolites; n=275,241 participants) and proteomic (n=2,905 proteins; n=50,575 participants) data from the UKB to generate two separate measures of dysregulation. Similar to the CLSA, dysregulation increased with age in both males and females whether quantified using metabolomics (p_male_<0.001 and p_female_=0.0086) or proteomics (p_male_=0.02 and p_female_<0.001) (Fig. 4a), and exhibited relatively modest heritability (metabolomics, h=3.2%, 95% CI 2.5%-3.8%, p<0.001; proteomics, h=4.4%, 95% CI -1.4%-10.3%, p=0.06). Furthermore, risk factors of poor health were broadly associated with higher dysregulation (Fig. 4b). Consistent with the CLSA, dysregulation significantly associated with survival, more so for males than females (hazard ratios (HR) = 1.08 to 1.28), as well as the incidence of most chronic conditions considered, more so when quantified using proteomics (HR= 1.06-2.04) as compared to metabolomics (HR=1.02-1.30) (Fig. 4d). Collectively, these results demonstrate that our Wasserstein distance-based approach can be used with different types of omics data from different cohorts to quantify dysregulation.

**Fig. 4.**
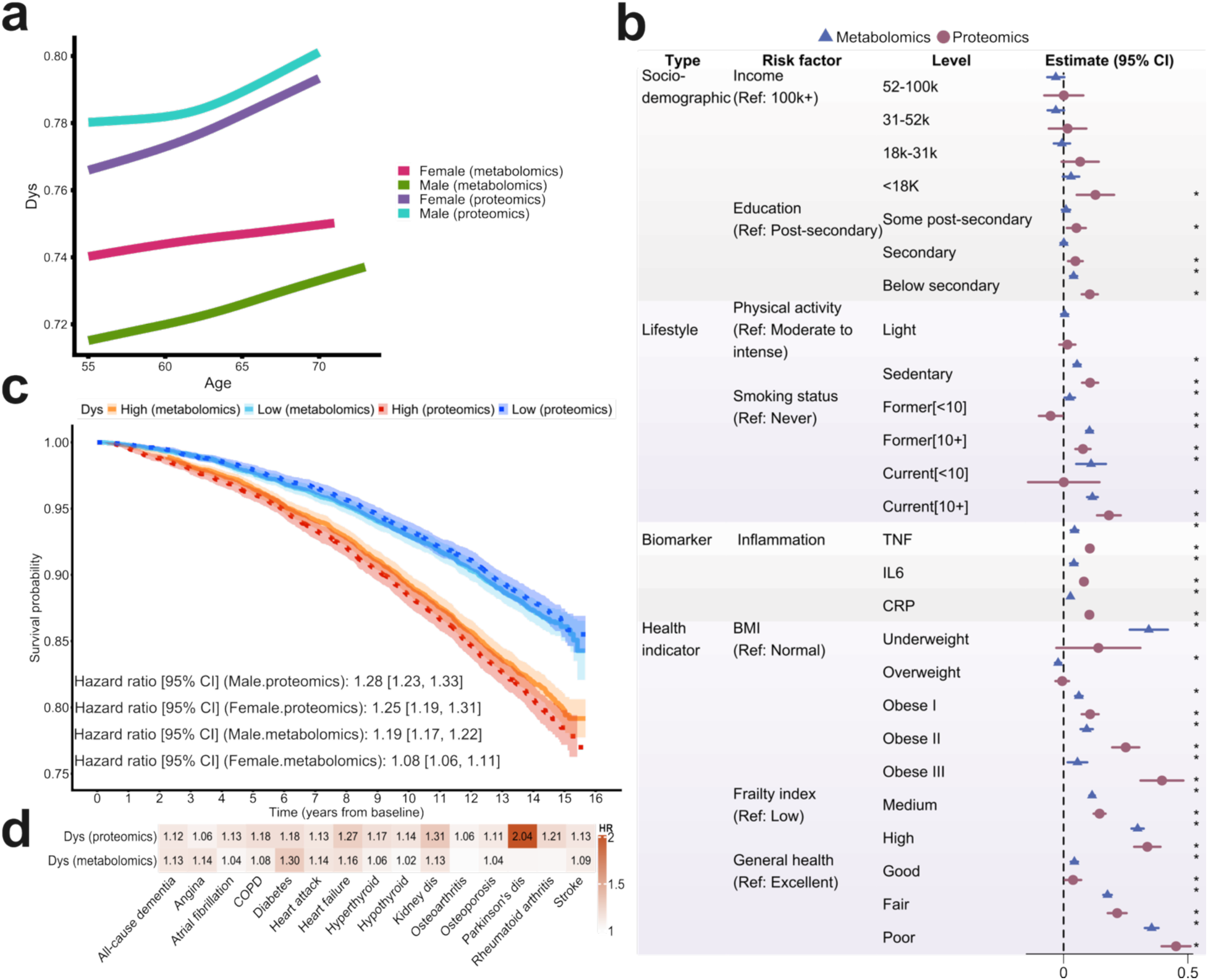
External validity of dysregulation associations in participants from the UK Biobank cohort. Biological dysregulation (Dys) quantified using either NMR metabolomics (n=159) or proteomics (n=2,905) data was correlated with **a,** age, separately for males and females. **b,** Associations of dysregulation with health risk factors and traits were obtained by linear regression, adjusting for age and sex, and reported as the standardized estimate and 95% confidence interval; Significance denoted by asterisks, p<0.05. **c,** Kaplan-Meier curves of dysregulation discretized into low and high groups based on tertiles. Cox proportional hazards models adjusted for age and the interaction between sex and dysregulation were used to estimate the 16-year mortality risk for males and females, reported as hazard ratios (HRs) with 95% confidence intervals. **d,** Associations with disease incidence reported as the HR (95% CI) from Cox proportional hazard models adjusted for age and sex. Only significant HRs (p<0.05) are displayed.

### Clustering metabolites and identifying biological systems

To explore the underlying systems structure of dysregulation, we clustered metabolites using an unsupervised method, Weighted Gene Co-expression Network Analysis (WGCNA), resulting in six groups of various sizes no smaller than 45 molecules (Fig. 5a). Systems were then characterized by correlating participant health traits with system principal component scores, which serve as composite abundance levels across assigned metabolites, and performing metabolite pathway enrichment analysis with MetaboAnalyst (Fig. 5b, 5c; Supplementary Tables 3). System zero (n=251 metabolites) comprised weakly correlated metabolites that could not cluster with other systems and was weakly associated with health traits and metabolic pathways (Fig. 5c). System one (n=229) correlated with health traits reflective of frailty (i.e. weak grip strength, low FVC, and a greater number of medications), multimorbidity, systemic inflammation (i.e. plasma TNF and monocyte count), and metabolic pathways, including amino acid metabolism, glucose homeostasis, and tryptophan catabolism. System two (n=109) was related to musculoskeletal traits (osteoporosis, back pain, weak grip strength, being underweight, and a low waist-to-hip ratio), cardiopulmonary traits (low FVC, low RBCs and oxygen transport), and lipid and fatty acid synthesis. System three (n=109) was related to cholesterol and sphingolipid metabolism. System four (n=68) was related to the high-load musculoskeletal system (i.e. strong grip strength, obesity, and high waist to hip ratio). Lastly, system five (n=45) was related to smoking, caffeine consumption, and pathways related to nicotine metabolism, caffeine metabolism, and flavan metabolism.

**Fig. 5.**
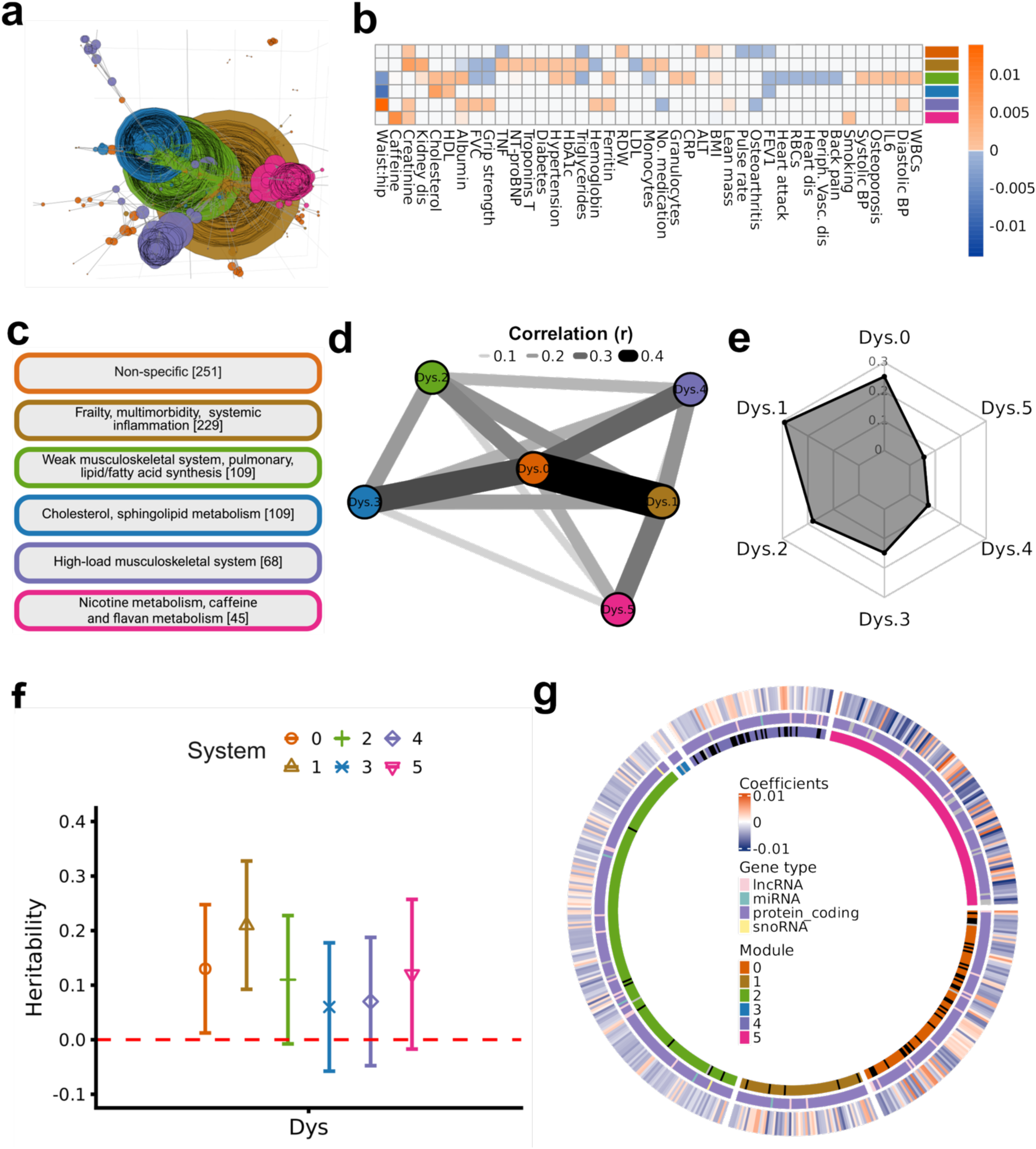
Investigating the heterogeneity of dysregulation across biological systems. **a,** Correlation network of biological systems (denoted by colour) identified using Weighted Gene Co-expression Network Analysis (WGCNA) of plasma metabolites. **b,** Associations of biological system principal component scores and health risk factors and indicators using Lasso regression, adjusted for age and sex. **c,** Characteristics of the systems identified by WGCNA and their relevance to phenotypes and pathways. The number of metabolites in each system is provided in brackets **d,** Correlation network of biological dysregulation quantified within each biological system, with Pearson correlation coefficients (r) denoted by edge shade and thickness. **e,** Spider plot showing the proportion of overall dysregulation attributed to dysregulation within each system. **f,** SNP-based heritability estimates for dysregulation in biological systems, with 95% confidence intervals; overlap with the red, dashed line denotes non-significance. **g,** Epigenome-wide association analysis of dysregulation in biological systems. Significant CpG sites are shown in the outer circle, with colors indicating the direction of association with dysregulation. The middle circle indicates the nearest-gene type for the CpGs, and the inner circle shows the biological systems attributed to the CpG sites. Grey segments in the middle circle represent CpGs with no specific gene type. Black segments in the inner circle indicate CpG sites that are shared across multiple biological systems.

### Characterizing the heterogeneity of system-specific dysregulation

To characterize the degree to which dysregulation is distinct and heterogenous across subsystems, we investigated pairwise correlation (Fig. 5d). We found that dysregulation was positively correlated across subsystems (all p<0.001), although unequally and of modest strength, where most correlations were between 0.1 and 0.3. System zero tended to be most strongly associated with other systems, particularly system one (r=0.48, p<0.001) and three (r=0.34, p<0.001). Dysregulation within each system correlated positively with overall dysregulation, although the magnitude varied substantially, ranging from 0.34 and 0.75 (all p<0.001) (Extended Data Fig. 3). None of the system-specific dysregulation fully explained the variation in overall dysregulation (Fig. 5e); System one accounted for the highest proportion of variation (0.29), while system five accounted for the lowest (0.05). Heritability of subsystem dysregulation varied between 6% and 21%, although only for systems zero, and one was it statistically significant. (Fig. 5f). Finally, we performed epigenome-wide association study (EWAS) in n=1,446 participants with available DNA methylation data to further characterize heterogeneity in our identified subsystems (Fig. 5g). Variation in a total of 566 CpGs were found to be significantly associated with system-specific dysregulation, most notably system two (n=196), five (n=123) and zero (n=107). Most CpGs were associated with protein coding genes, and downregulation was observed at 388 CpG sites, with significant enrichment of downregulated CpGs in system two (n=149, FDR=0.03). Importantly, very few CpGs (n = 32) were associated with multiple systems, suggesting unique underlying epigenetic mechanisms of dysregulation across systems. Overall, the identified systems exhibited different characteristics and were involved in distinct biological processes.

### Risk factors of system-specific dysregulation

Associations between risk factors and dysregulation across different biological systems were quite similar, although some notable differences were observed (Fig. 6 and Extended Data Fig. 4). For example, the effect of smoking status was substantially higher in system five (i.e. current, heavy smokers; d=1.02), while being underweight was more strongly associated with higher dysregulation in systems two (d=0.69) and five (d=0.55), and severe obesity with system one (d=0.57). High FI was associated with higher dysregulation in all systems, with the strongest effects related to systems one (d=0.55).

**Fig. 6.**
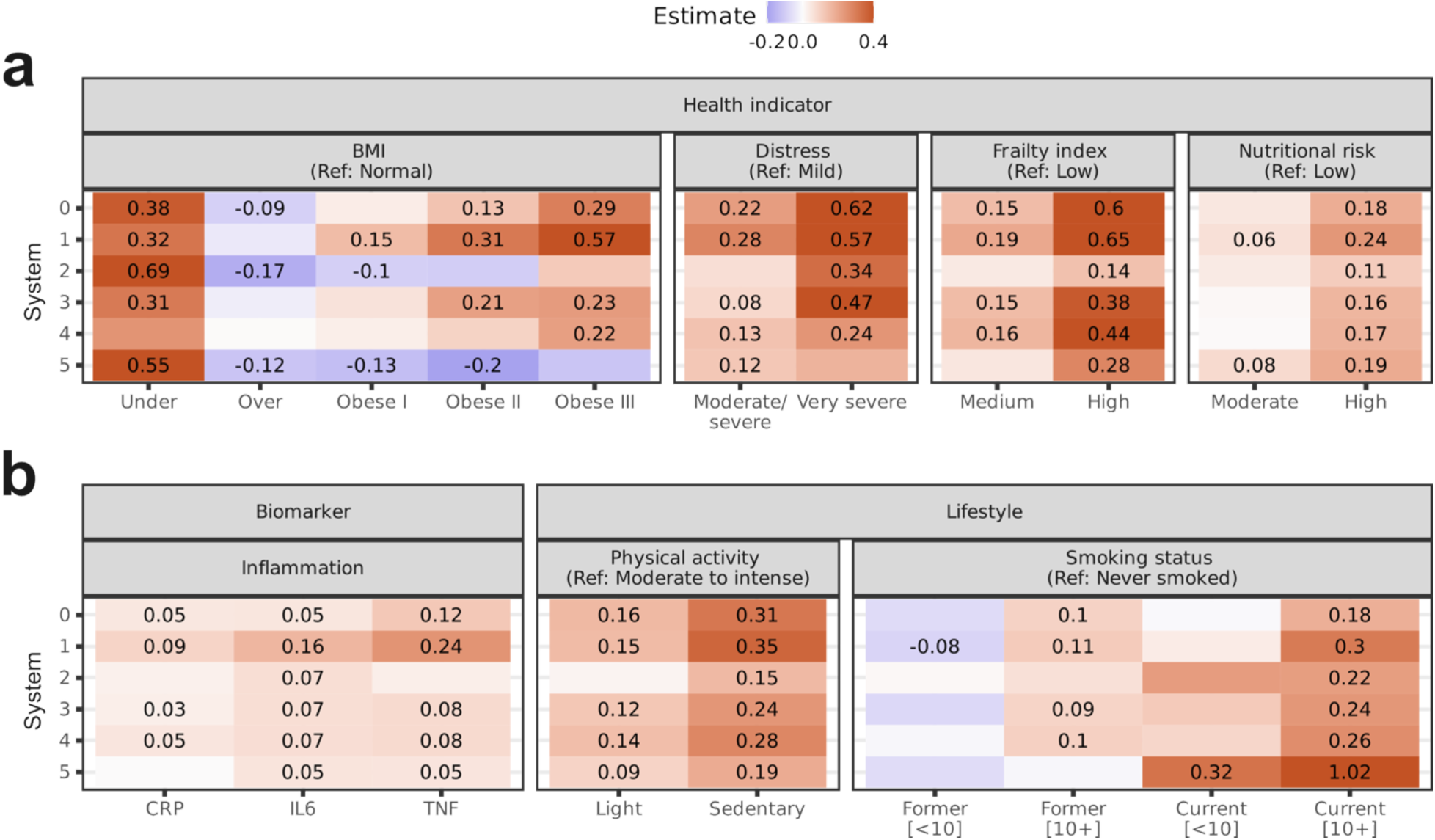
Risk factors of dysregulation within biological systems. Associations of system-specific dysregulation with risk factors, including **a,** health indicators, and **b,** biomarker and lifestyle factors, estimated using linear regression, adjusted for age and sex. Standardized regression coefficients for estimates determined to be statistically significant (p<0.05) are shown in each respective cell.

### System-specific dysregulation predicts adverse outcomes heterogeneously

We further assessed system-specific dysregulation as predictors of outcomes, including incidence of chronic diseases, mortality risk, cognitive decline, physical function decline, and healthcare utilization (Fig. 7 and Extended Data Fig. 5). Significant heterogeneity was observed in the associations of system-specific dysregulation with incident chronic conditions over six years (Fig. 7a). For example, only system two was associated with the incidence of dementia and Alzheimer’s disease (AD, 41 cases; HR=1.68, p<0.001) and hyperthyroidism (51 cases, HR=1.34, p=0.034). In contrast, dysregulation in system one was associated with the incidence of several diseases, the most significant being kidney disease (156 cases, HR=1.86, p<0.001), but remained the only system associated with hypothyroidism (278 cases, HR=1.22, p=0.001) and rheumatoid arthritis (RA, 156 cases; HR=1.25, p=0.01). Additionally, we sought to assess the independent contribution of dysregulation in each system to the risk of incident diseases by accounting for dysregulation in other systems (Extended Data Fig. 5c). Notably, heart disease, hypothyroidism, and RA were mainly attributed to dysregulation in system one, whereas dementia and AD were mainly associated with system two.

**Fig. 7.**
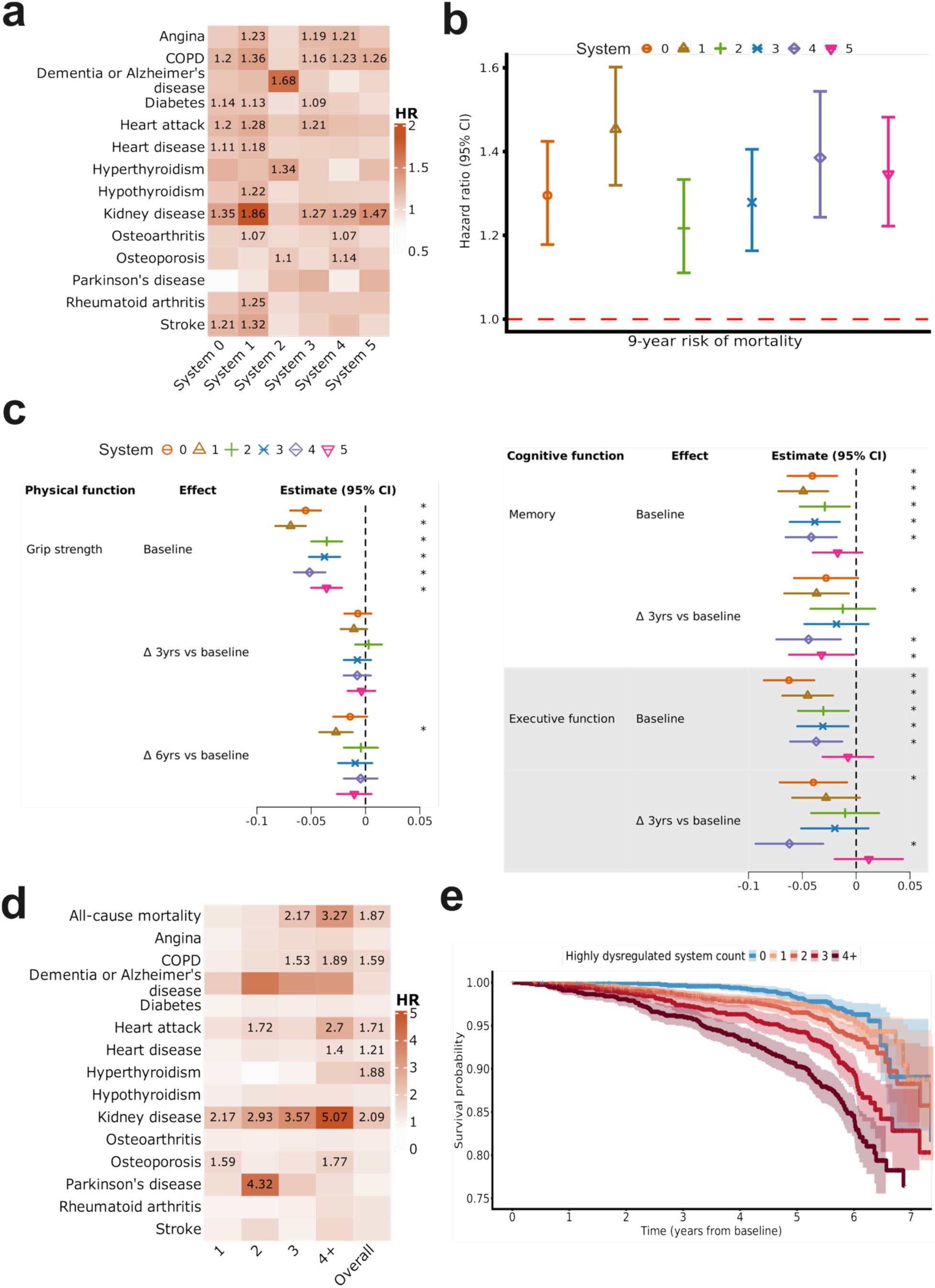
Outcomes associated with system-specific dysregulation and estimated cumulative effects. **a,** Associations with disease incidence using Cox proportional hazard regression, adjusting for age and sex, where only significant hazard ratios (HR;p<0.05) are displayed. **b,** Nine-year risk of mortality associated with system-specific dysregulation, by Cox proportional hazards regression, adjusted for age and sex, and HR and 95% confidence intervals. **c,** Standardized estimates and 95% confidence intervals reported for the association between system-specific dysregulation and grip strength, and memory and executive function. Δ 3yrs or Δ 6yrs vs. baseline refers to the 3-year or 6-year change relative to baseline, and significance denoted by asterisks (p<0.05). **d,** The age- and sex-adjusted hazard of all-cause mortality and disease incidence associated with high dysregulation in one, two, three, or four or more (4+) biological systems, as compared to no systems, and high vs. low dysregulation overall. Only significant HRs (p<0.05) are displayed. **e,** Kaplan-Meier analysis of all-cause mortality for participants exhibiting high dysregulation within none, one, two, three, and more than three (4+) systems. The x-axis is truncated at 7 years for clearer visualization.

All-cause mortality was significantly associated with dysregulation in all systems, although to varying magnitudes, the greatest being system one (HR [95% CI]= 1.45 [1.32, 1.6]) and the lowest being system two (1.22 [1.11, 1.33]) (Fig. 7b). Dysregulation in all systems except system two was associated with weaker grip strength (Fig. 7c, left) and slower walking speed (Extended Data Fig. 5a) at baseline with different magnitudes. However, only system one was associated with reduced grip strength over six years (d=-0.03). For cognitive function (Fig. 7c, right), dysregulation in all systems except system five was associated with lower performance in both memory and executive function tasks at baseline. Over time, dysregulation in system four was the strongest predictor of decline in both memory (d=-0.04) and executive function (d=-0.06), while dysregulation in systems one (d=-0.04) and five d=-0.03 predicted memory decline and, and dysregulation in system zero predicted decline in executive function (d=-0.04). All systems were broadly associated with emergency department visits or overnight hospitalization (OR=1.13-1.36) (Extended Data Fig. 5b).

### Cumulative health burden of multi-system high dysregulation

Although system-specific dysregulation was found to be broadly associated with numerous health traits and outcomes, it remains unclear whether they merely serve as proxies for an overarching dysregulation phenomenon, or if they operate synergistically. To investigate this, we estimated the risk of mortality and incident disease related to a cumulative score representing the number of systems for which a participant was within the upper tertile (i.e., high) of dysregulation and compared it to being in the upper tertile for our overall dysregulation measure. We identified n=1,205 individuals (16%) who did not exhibit high dysregulation in any system, while n=1,959 (26%), 1,830 (24%), 1,238 (17%), and 1,239 (17%) individuals exhibited high dysregulation in one, two, three, and at least four systems, respectively. In contrast, n=2,540 individuals exhibited high overall dysregulation. Interestingly, higher cumulative dysregulation was associated with an increased risk of poor health outcomes, more so than exhibiting high overall dysregulation; for example, cumulative high dysregulation burden in 3 (HR=2.17, p<0.001) or 4+ (HR=3.27, p<0.001) systems was associated with a higher risk of mortality than overall high dysregulation (HR=1.87, p<0.001) (Fig. 7d, 7e). The cumulative burden of system-specific dysregulation was also more strongly associated with many incident diseases. In particular, high dysregulation in 2 (HR=2.93, p=0.003), 3 (HR=3.57, p<0.001), or 4+ (HR=5.07, p<0.001) systems were more strongly associated with an increasing risk of kidney disease than overall high dysregulation (HR=2.09, p<0.001) (Fig. 7d). Hence, compared with overall high dysregulation across the whole body, multi-system high dysregulation showed stronger associations with adverse outcomes, suggesting that multi-system high dysregulation can be disproportionately detrimental, which cannot be captured by overall high dysregulation.

## Discussion

We used plasma metabolites to study biological dysregulation as they are the mediators and end products of cellular processes, making them uniquely reflective of disease or physiological changes^48^, whether measured in the circulation or in peripheral tissues^49,50^. Our novel measure of biological dysregulation based on Wasserstein distance is computationally efficient and does not suffer from issues related to collinearity of biomarkers. Using this method, we found biological dysregulation to be associated with both risk factors and health outcomes, which we validated in data from the UK Biobank. In that cohort, we showed our method could be applied to both metabolomic as well as proteomic data, with dysregulation in either being broadly associated with health traits and outcomes. We also assessed sex differences across a variety of risk factors and outcomes. In particular, we observed that high frailty, a sedentary lifestyle, and higher levels of inflammatory markers were more strongly associated with dysregulation in males than in females, as was reduced grip strength over time. The risk of mortality associated with dysregulation was also higher in males but was not statistically significant. These findings support an inherent opposing relationship between biological dysregulation and resilience, whereby women more effectively respond to stressors than men, and maintain lower levels of dysregulation as a consequence. Arbeev and colleagues^51^ observed similar trends in their analysis of dysregulation trajectories, concluding that males generally have lower resistance to stressors and a greater risk of mortality with age in relation to dysregulation. However, Cohen and colleagues^9^ failed to find sex differences in mortality risk attributable to dysregulation. Given the inconsistencies among these studies with respect to dysregulation quantification approaches, sample populations, and statistical power, the discrepancies in findings may not be surprising and point to a greater need for homogeneity in study designs moving forward.

The homeostatic and allostatic processes that are critical in determining the resilience of an organism depend on a complex, overlapping network of fundamental biological systems^8^. While previous studies have attempted to characterize these biological systems, mostly using literature-driven manual curation or domain mapping approaches^8–12^, they featured some notable limitations. First, they were generally limited to relatively smaller sets of markers. Given the significant overlap in biomarkers related to different organs or physiological processes, a limited set may not be adequate to capture the interconnectedness of resultant systems and support robust inference^8^. Second, reliance on biomarker annotation to derive systems is problematic since many markers will be annotated to more than one organ or physiological process, markers that have been investigated more intensively will likely be annotated better^52^, and annotation may be tissue-specific and therefore fail to reflect the biological context elsewhere in the body^53^. Here, we used a dense panel comprising 811 metabolites from different super pathways, such as lipids, amino acids, xenobiotics, and cofactors and vitamins, and Weighted Gene Co-expression Network Analysis (WGCNA)^36^, an unsupervised, data-driven method, to identify systems of co-regulated metabolites without relying on predefined pathway annotations. Although this resulted in systems without definitive classification regarding biological or physiological processes, the topological approach of WGCNA ensures that they represent functionally coherent networks of densely connected metabolites in a mutually exclusive manner^54^. Even though these systems were distinct regarding the metabolites assigned, we nonetheless observed substantial overlap in the association between composite abundance levels (i.e. system principal component scores) and health traits and indicators, both in concordant and opposing directions. For example, system one, which was associated with amino acid metabolism and transport, was positively correlated with triglyceride and monocytes, as shown in previous studies^55,56^. However, system two, which was associated with fatty acid metabolism, was negatively correlated with both indicators, as shown previously^57^. In contrast, systems one and two were both positively associated with glycated hemoglobin (i.e. HbA1c), potentially through their respective involvement in the ‘Glucose Homeostasis’ and ‘Synthesis, Secretion, and Inactivation of Glucagon-like Peptide-1 (GLP-1)’ pathways, as shown in previous studies^58,59^. Given this overlap, we hypothesized that dysregulation would be strong between systems; however, mostly weak, yet statistically significant, positive correlations (r<0.3) were observed, although all of which were statistically significant and positive. This observation suggests that dysregulation across systems is likely coordinated, or at least evolves in a parallel fashion, which was also concluded in the work of Li et al^8^. An exception to this observation was dysregulation in system zero, which exhibited moderate correlations (0.15<r<0.48) with other systems, even though the metabolites in system zero were those that failed to exhibit marked correlation with others and therefore could not be clustered. Nonetheless, dysregulation in system zero was broadly associated with health risk factors and outcomes, and was found to exhibit significant heritability. It remains unclear whether the unassigned metabolites in system zero are noise or redundancy from other systems.

Another significant finding from our work was that dysregulation across systems exhibited significant heterogeneity with regards to associations with a diverse array of health risk factors, traits, and outcomes. This echoes previous theoretical views^60^ that these systems likely overlap in supporting the biological mechanisms underlying resilience. Although this overlap mainly manifested as associations with health traits and outcomes in a similar directionality, there were instances of marked discordance. Notably, dysregulation within system one was more strongly associated with obesity than dysregulation in other systems, while dysregulation within system two was significantly associated with being underweight at nearly the same magnitude. Interestingly, system two was also found to be enriched with metabolites related to the pathway ‘synthesis, secretion, and inactivation of Glucagon-like Peptide-1 (GLP-1)’, which is known to promote weight loss mainly by targeting the brain to decrease appetite and the stomach to slow the digestive process^61^. Similar to health risk factors and traits, there was a great deal of concordance in the associations amongst system-specific dysregulation and health outcomes, although some striking differences were apparent. For example, dysregulation in system two, which was enriched for metabolites related to fatty acid metabolism, including pathways such as mitochondrial beta-oxidation, fatty acid synthesis, and adipogenesis, was uniquely associated with the incidence of dementia and AD, hyperthyroidism, and osteoporosis, although the underlying mechanisms are yet to be identified. While fatty acids are known to play a myriad of roles in the regulation cellular function, authors commonly point to evidence indicating changes in fatty acid composition as being a critical regulator of immune and inflammatory activation, dysfunction of which driving disease incidence and worsening pathology^62^. In contrast, some outcomes were found to be commonly associated with a number of systems, namely chronic obstructive pulmonary disease (COPD) and kidney disease, and to a lesser extent, angina, diabetes, and heart attack. Given that these conditions often coexist, in particular COPD with kidney disease^63^ and diabetes with heart attack^64^, our observed overlap in outcome associations likely reflects the intertwined nature of the systems identified and suggests they represent fundamental pathways and processes that broadly determine overall health and link conditions into a network that Barabási and colleagues coined as a “diseasome”^65^. Intuitively, this suggests that the effect of multi-system dysregulation would be cumulative, where the risk of adverse health outcomes increases along with the number of systems that are in a dysregulated state. Indeed, our data supports this hypothesis. For example, high dysregulation in more than three systems was associated with more than double the risk of kidney disease as compared to high overall dysregulation, while the risk of all-cause mortality was nearly 75% greater. These findings support the presence of a non-additive and potentially synergistic health burden associated with multi-system dysregulation and suggest that interventions targeting multiple systems may be more effective in preventing adverse health outcomes than interventions targeting a single system in isolation^66^.

To the best of our knowledge, this is one of the most comprehensive studies to date examining biological dysregulation in older adults and characterizing system-specific dysregulation through a network approach using a wide range of health traits. Major strengths include a large, well-balanced sample that is generalizable to Canadian adults aged 45 to 86, comprehensive phenotyping with up to nine years of follow-up, and external validation in a separate cohort. However, there are some limitations that are important to consider. First, our findings are limited to plasma metabolites, which may not accurately reflect the systems structure of dysregulation in peripheral tissues. Second, although we validated our novel measure of dysregulation in another large population-level cohort, our findings remain mostly reflective of Caucasian individuals; hence, replication of our findings in cohorts with different predominant ethnicities is needed. Third, the CLSA population includes participants aged 45 years and older, whereas the first phase of molecular and structural reprogramming due to aging has been suggested to occur around 33 years old^67^. Therefore, our results may not be generalizable to younger populations. Finally, nearly half of metabolites we studied have no annotation, limiting our ability to properly identify the pathways of each system.

In summary, this study provides a network view of biological dysregulation based on hundreds of metabolites across biological systems and explores sex differences in whole-body dysregulation with health risk factors and outcomes. Through the use of Wasserstein distance, a simple, computationally efficient statistical measure applicable to low- and high-dimensional biomarker data, and across different cohorts, we show that dysregulation partially overlaps across multiple biological systems. This suggests that breakdown in one system may result in breakdown and dysregulation in other interdependent systems, resulting in a cumulative burden on the risk of mortality and morbidity. Given its association with several modifiable risk factors, including physical activity, smoking, nutrition, and BMI, our findings suggest that lifestyle and dietary interventions could be used to target dysregulation and thereby promote healthy aging. Although we used the full metabolite panel to have a comprehensive view of system-level dysregulation, identifying a parsimonious subset of metabolites that achieves similar results may improve clinical applicability while reducing costs.

---

**Extended Data Fig. 1.**
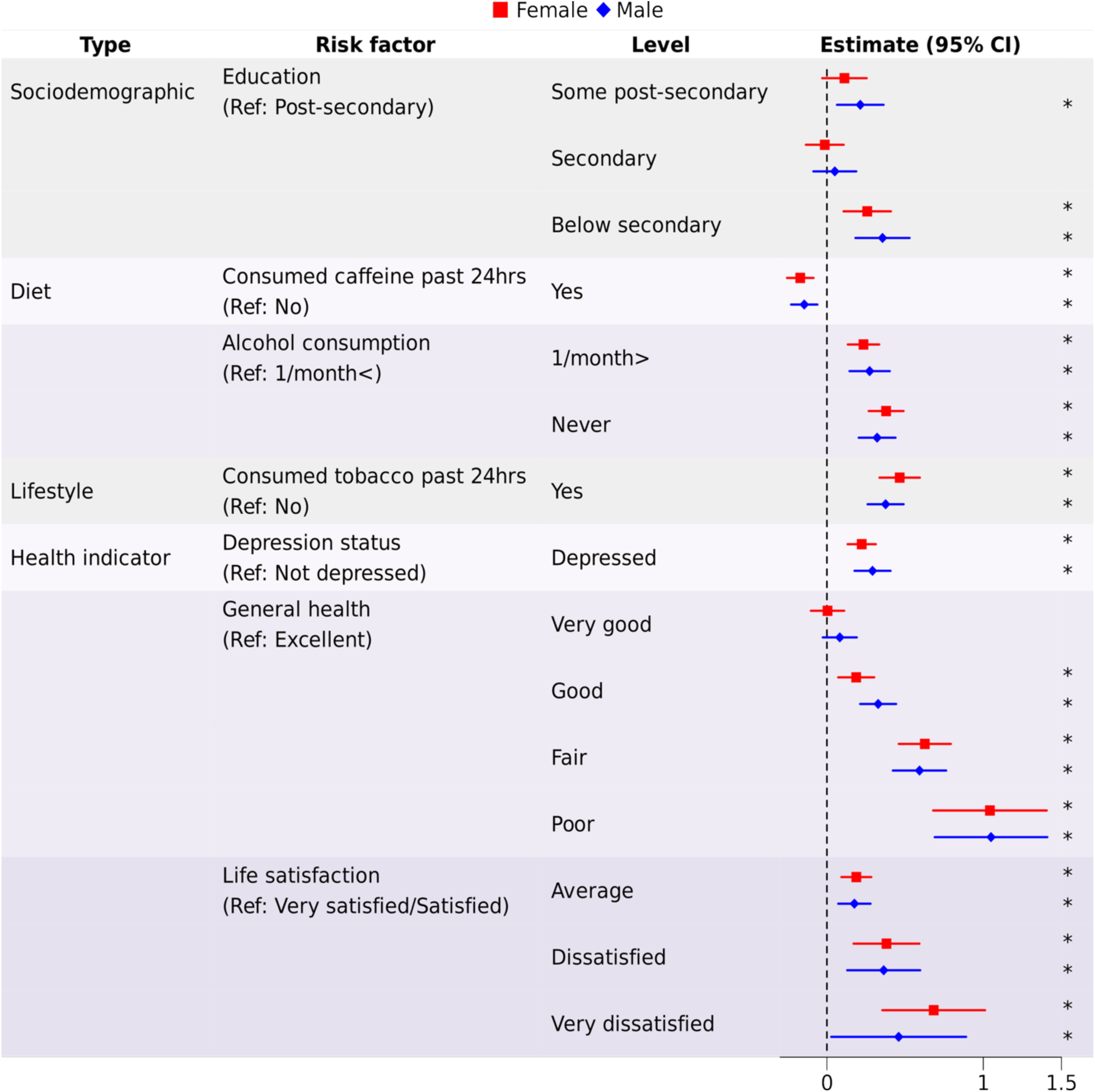
Sex-specific associations of dysregulation with health risk factors. Standardized estimates and 95% confidence intervals from linear regression analyses describing the association between dysregulation and sociodemographic, diet, lifestyle factors, and health indicators. All models include the fixed effects of age and the interaction between sex and a single predictor. The dashed line indicates no correlation, while the asterisks ‘*’ indicates p<0.05.

**Extended Data Fig. 2.**
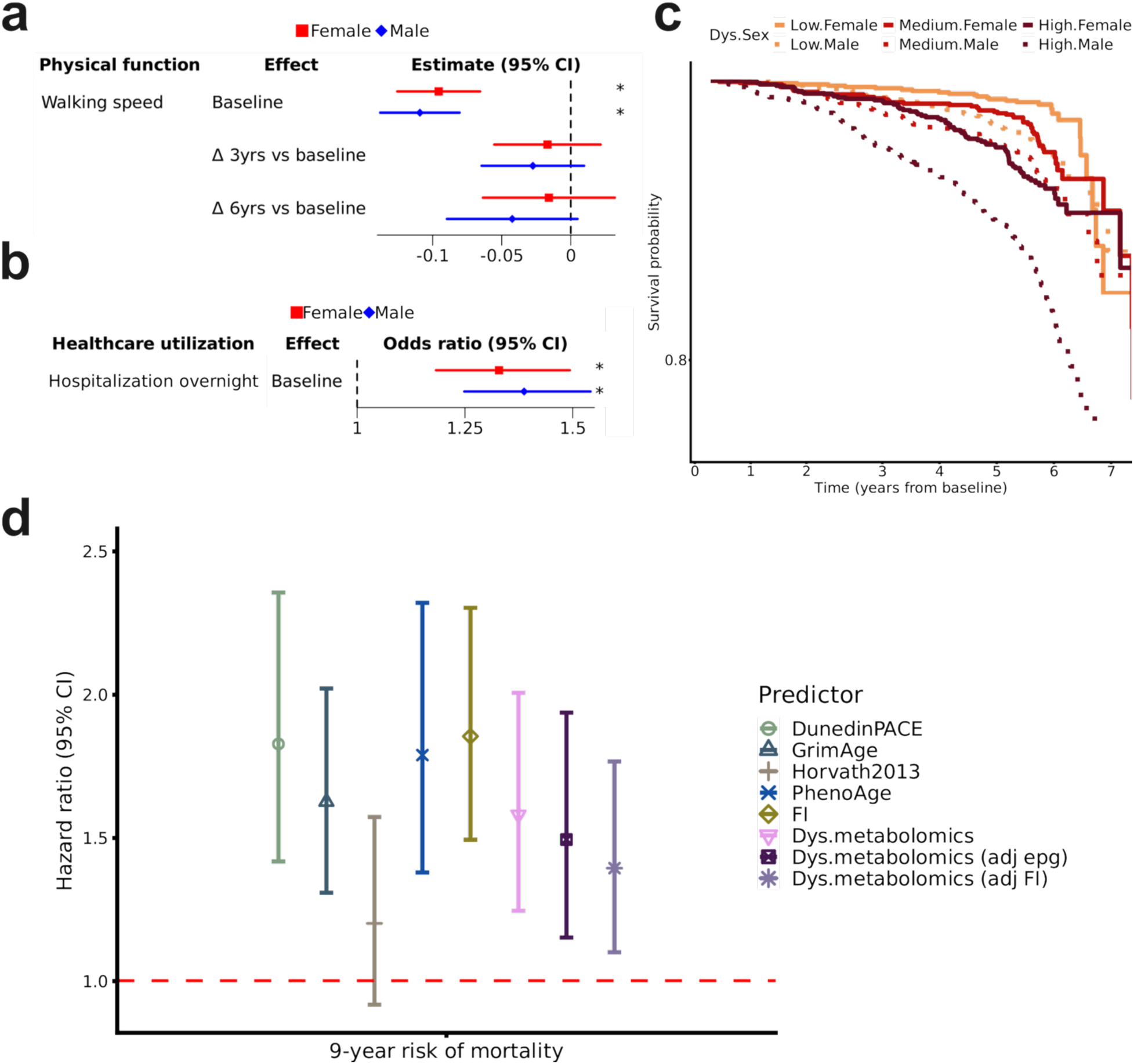
Sex-specific associations of dysregulation with outcomes and comparative survival analysis with epigenetic aging clocks and frailty index (FI). **a,** Standardized estimates and 95% confidence intervals reported for the association between dysregulation and walking speed. Δ 3yrs or Δ 6yrs vs. baseline refers to the 3-year or 6-year change relative to baseline, and significance denoted by asterisks (p<0.05). **b,** Odds ratios and 95% confidence intervals for overnight hospitalization associated with dysregulation. **c,** Associations with all-cause mortality indicated by Kaplan-Meier curves, with dysregulation tertilized as low, medium or high for males and females. The x-axis is truncated at 7 years for clearer visualization. **d,** Nine-year mortality risk associated with epigenetic aging clocks, frailty index (FI), and dysregulation (Dys) based on metabolomics with and without adjustment for epigenetic aging clocks (epg) or FI). Cox proportional hazard models adjusted for age and sex were used to estimate hazard ratios with 95% confidence intervals.

**Extended Data Fig. 3.**
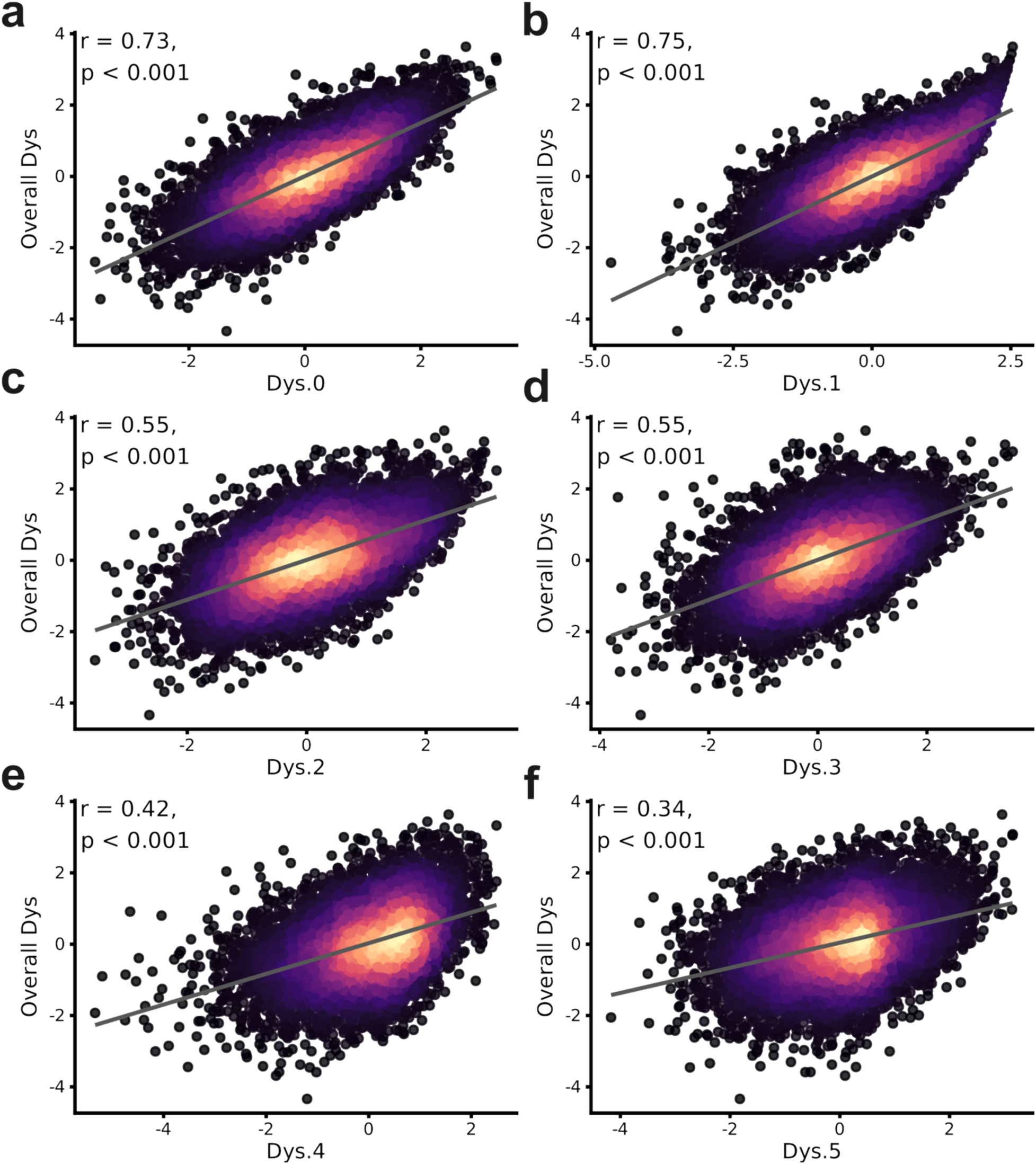
Correlations between overall dysregulation (Dys) and system-specific dysregulation (a-f). Pearson’s correlation (r) between overall dysregulation (y-axis) and system-specific dysregulation (x-axis), along with the corresponding p-value, is shown.

**Extended Data Fig. 4.**
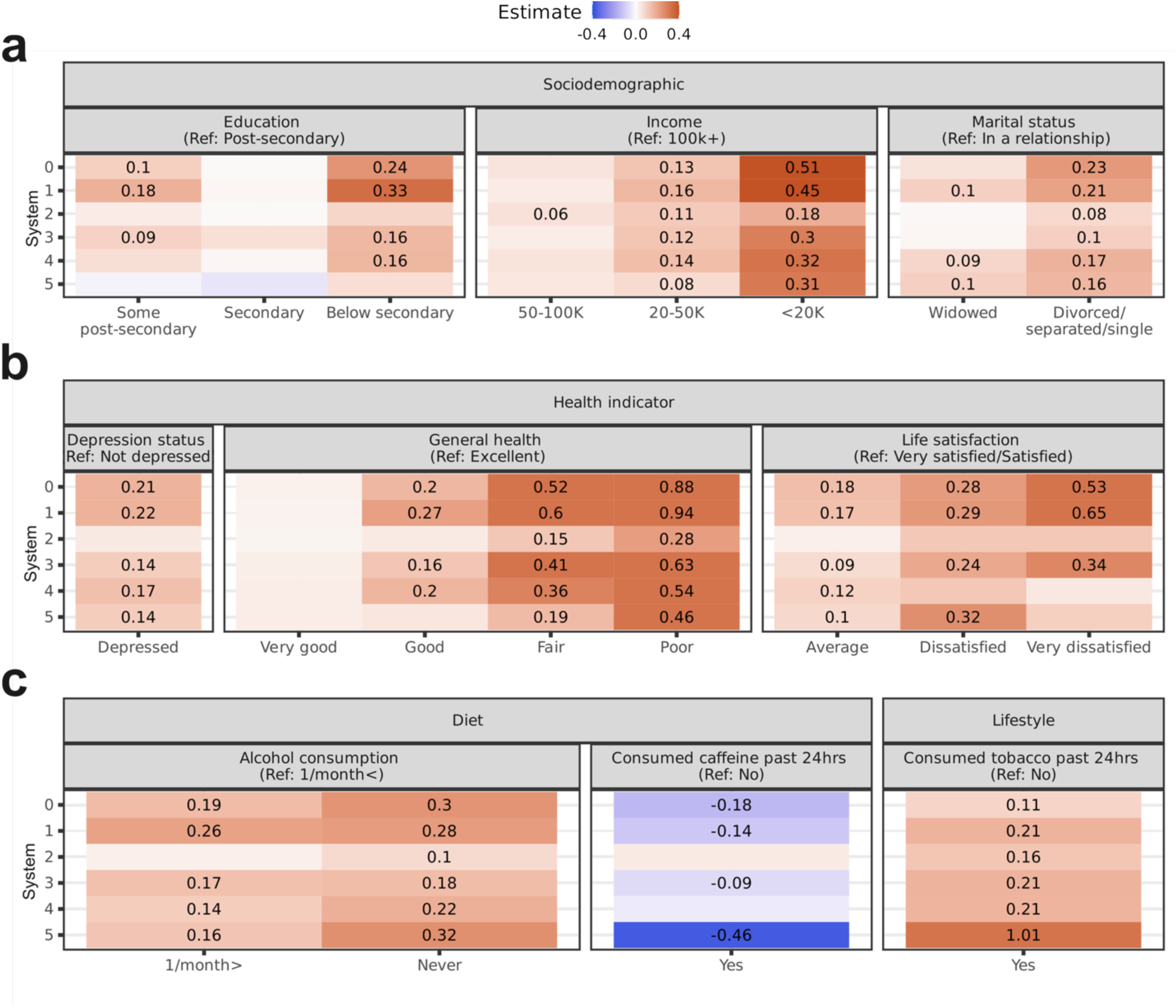
Risk factors of system-specific dysregulation. Associations of system-specific dysregulation with risk factors, including **a,** sociodemographic factors **b,** health indicators, and **c,** diet and lifestyle factors, estimated using linear regression, adjusted for age and sex. Standardized regression coefficients for estimates determined to be statistically significant (p<0.05) are shown in each respective cell.

**Extended Data Fig. 5.**
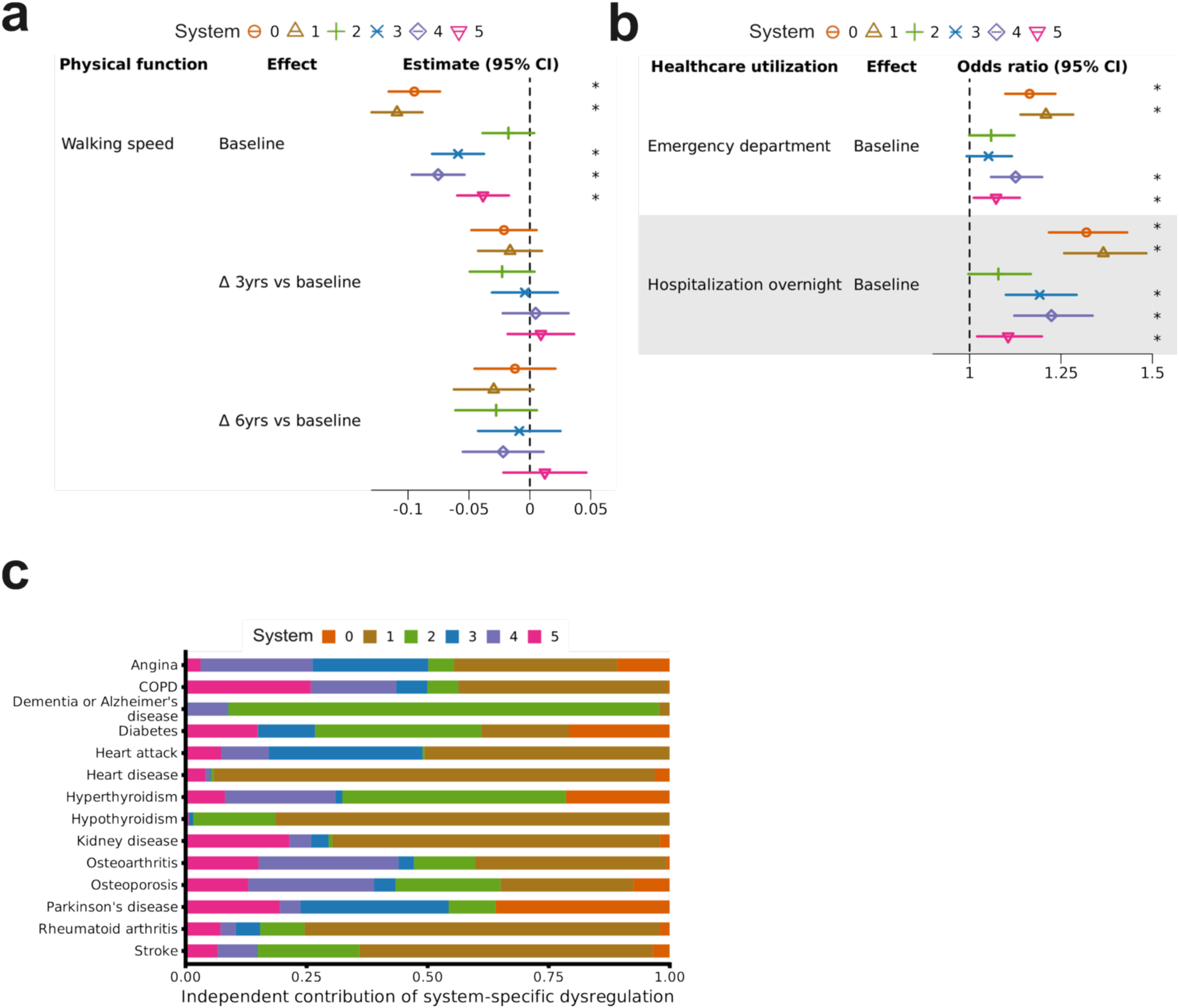
Outcomes associated with system-specific dysregulation. **a,** Standardized estimates and 95% confidence intervals reported for the association between system-specific dysregulation and walking speed. Δ 3yrs or Δ 6yrs vs. baseline refers to the 3-year or 6-year change relative to baseline, and significance denoted by asterisks (p<0.05). **b,** Odds ratios and 95% confidence intervals for emergency department visit and overnight hospitalization associated with dysregulation. **c,** Independent contributions of system-specific dysregulation to each disease incidence prediction. All regression models are adjusted for age and sex.

## Notes

### Competing Interest Statement

The authors have declared no competing interest.

### Funding

- Canadian Institutes of Health Research

